# Leveraging a WGS compression and indexing format with dynamic graph references to call structural variants

**DOI:** 10.1101/2020.04.24.060202

**Authors:** Adam C. English, Nils McCarthy, Robert Flickenger, Surabhi Maheshwari, Lisa Meed, Alindrina Mangubat, Sri Niranjan Shekar

## Abstract

We introduce a novel structural variant calling pipeline (BioGraph) that leverages a read compression and indexing format to discover alleles, assess their supporting coverage, and assign useful quality scores. To evaluate BioGraph’s performance, we run five sequencing replicates of the individual HG002 through the BioGraph variant calling pipeline, as well as two other short-read sv-calling pipelines. BioGraph detects the GIAB benchmark SVs at a peak sensitivity of ≈59% compared to ≈42% sensitivity from the other pipelines. The overall precision of BioGraph is lower than other pipelines (≈81% and ≈90%, respectively), however, adjusting for quality score, BioGraph calls were sensitive to a greater number of SV calls given the same false discovery rate compared to the other pipelines. Cumulatively ≈77% of GIAB benchmark SVs are discovered by BioGraph in at least one replicate. After merging discovered calls and running BioGraph Coverage to create a squared-off project-level VCF, we find ≈90% percent of discovered true positive alleles have at least 5x coverage in all replicates, thus increasing per-replicate recall of alleles having at least 1x coverage to ≈76.9%.

## Background

Most current next-generation sequencing secondary analysis pipelines map short-reads (≈150bp) to a reference genome. Since these reads can be confidently placed while spanning smaller SNP/INDEL (<≈50bp) variations [7], pileups of the bases over positions in the reference are measured. However, for larger changes, multiple signals are used from the short-read alignments to infer the presence of structural variation (SVs >≈50bp) such as soft-clipping, discordant read-pair distances, and abnormalities in coverage [16]. Many existing software packages leverage these signals to call SVs [4].

Since these tools attempt to elucidate different alignment signals to infer the presence of SVs, each tool typically varies in performance across particular type and size ranges of SVs.

This has led to the development of consensus calling pipelines that combine SVs reported by various tools into a single set of confident SV calls [20]. Many SV discovery tools do not report exact breakpoints or fully resolved sequences of alleles, creating complications for downstream applications such as for annotation and multi-sample merging.

When discovering SVs with short-read alignment based inference, two difficulties arise. First, short-read sequences are not long enough to disambiguate all genomic repeats [1, 3], which are an important mechanism in SV formation [5]. Second, SV callers may be susceptible to false positives from calling SVs using alignment information of reads where only part of the read maps to the reference or when using the expected mate pair distance, resulting in reference bias [18].

Assembly-based methods may be less susceptible to mapping artifacts [19] as they observe the relationship between reads as well as between the reads and the reference. Assembly based methods have shown relatively higher sensitivity to SV detection [10, 17]. However, de novo assembly of short-reads requires greater compute resources than read alignment and often special library prep [13, 15].

In addition to discovering SVs, research into confirming the presence of alleles and genotyping them within samples is ongoing [6]. Methods leveraging graph-based representations of known variation alongside a reference genome have begun to show promise for genotyping SVs [8].

We introduce a method of storing and accessing short-read next generation sequencing data (BioGraph Format) that allows for rapid querying of reads by sequence. We leverage this format with our BioGraph Discovery software to call variants using reference-guided read overlap assembly. Taking the resulting VCF with sequence resolved calls, we again leverage the BioGraph Format with our BioGraph Coverage software to quickly assess coverage supporting these putative alleles. We describe a machine learning model, BioGraph QUALclassifier, that uses coverage signatures to assign quality scores to discovered alleles to increase specificity and assist prioritization of variants. We benchmark five technical replicates of an individual (HG002) against the Genome in a Bottle Tier 1 SV set [21] alongside other SV detection pipelines to illustrate relative performance. Finally, we merge discovered calls and confirm the presence of alleles with BioGraph Coverage to assess the per-replicate increase in recall over discovery.

## Results

This work used five replicates of the Personal Genome Project’s Ashkenazi Jewish Proband HG002. Replicate 0 contains approximately 29.5x coverage and the other four range between 44.3x and 48.9x coverage; all had 150bp reads. Full details of the sequencing used can be found in the Supplementary 1.

The five sequencing replicates were run through the BioGraph variant calling pipeline, Manta, and Parliament2. Results were compared to the Genome in a Bottle v0.6 Tier 1 SV set using Truvari (Table 1). The average sensitivity of the BioGraph pipeline is ≈59% compared to ≈43% and 38% for Manta and Parliament2 respectively. The overall precision of BioGraph is lower (81%) than other callers (≈91% Manta, and ≈89% Parliament2). The point estimate of the average F1-score of BioGraph across replicates is higher than the other pipelines by approximately 10pp.

**Table 1.** Average SV (>=50bp) performance ± 1 standard deviation (across 5 replicates) between BioGraph, Manta, and Parliament2. BioGraph and Manta were compared to the truth set using Truvari matching parameters which require alternate allele sequence similarity >= 70% whereas Parliament2 was not required to pass this threshold since it does not report sequence resolved calls. Details of every replicate’s performance can be found in Supplementary 2

|  | BioGraph | Manta | Parliament2 |
| --- | --- | --- | --- |
| **True Positives** | 5,659 ± 303 | 4,129 ± 354 | 3,656 ± 285 |
| **False Negatives** | 3,982 ± 303 | 5,512 ± 354 | 5,985 ± 285 |
| **False Positives** | 1,449 ± 75 | 402 ± 29 | 469 ± 12 |
| **Precision** | 81% ± 1.23% | 91% ± 0.50% | 89% ± 0.93% |
| **Recall** | 59% ± 3.15% | 43% ± 3.67% | 38% ± 2.96% |
| **F1** | 68% ± 2.49% | 58% ± 3.57% | 53% ± 3.13% |

In addition to assessing the total number of alleles discovered, we observe how the tools perform across type/size categories on average across the replicates (Table 2). The recall suggests that the sensitivity to SVs with the BioGraph pipeline was higher on average across the replicates for variants between 50 and 1000 base pairs for both insertions and deletions.

**Table 2.** Recall of Deletions (DEL) and Insertions (INS) across four size bins. The first number in each cell is the total number of TPs called. The percent in parenthesis is the ratio of TPs called over the number of TPs in the truth-set for that variant type and within that size range. *Parliament2 does not require sequence similarity to match calls to the truth-set. **Manta reports INS over 1kbp, but does not report the associated sequence, and therefore has no TPs for that cell. Details for each replicate can be found in Supplementary 3.

| SVType | SVSize | BioGraph | Manta | Parliament2* |
| --- | --- | --- | --- | --- |
| DEL | [50, 100) | 1,098 (76%) | 853 (59%) | 566 (39%) |
|  | [100, 300) | 631 (65%) | 512 (52%) | 174 (18%) |
|  | [300, 1000) | 1,011 (82%) | 989 (80%) | 996 (80%) |
|  | >= 1,000 | 375 (71%) | 439 (83%) | 486 (92%) |
| INS | [50, 100) | 1,004 (67%) | 661 (44%) | 682 (45%) |
|  | [100, 300) | 657 (48%) | 361 (26%) | 371 (27%) |
|  | [300, 1000) | 834 (46%) | 314 (17%) | 315 (17%) |
|  | >= 1,000 | 48 (6%) | 0 (0%)** | 89 (12%) |

Another important metric for benchmarking is the consistency with which pipelines call the same TPs (true positives) between replicates. Figure 1 shows the counts of how many replicates in which each allele was found after uniting all the calls. True positives were united by the GIAB call to which they matched. To unite false positive calls, a VariantKey [2] was assigned to each variant based on its chromosome, position, and reference/alternate alleles’ sequence in order to merge between replicates.

**Figure 1.**
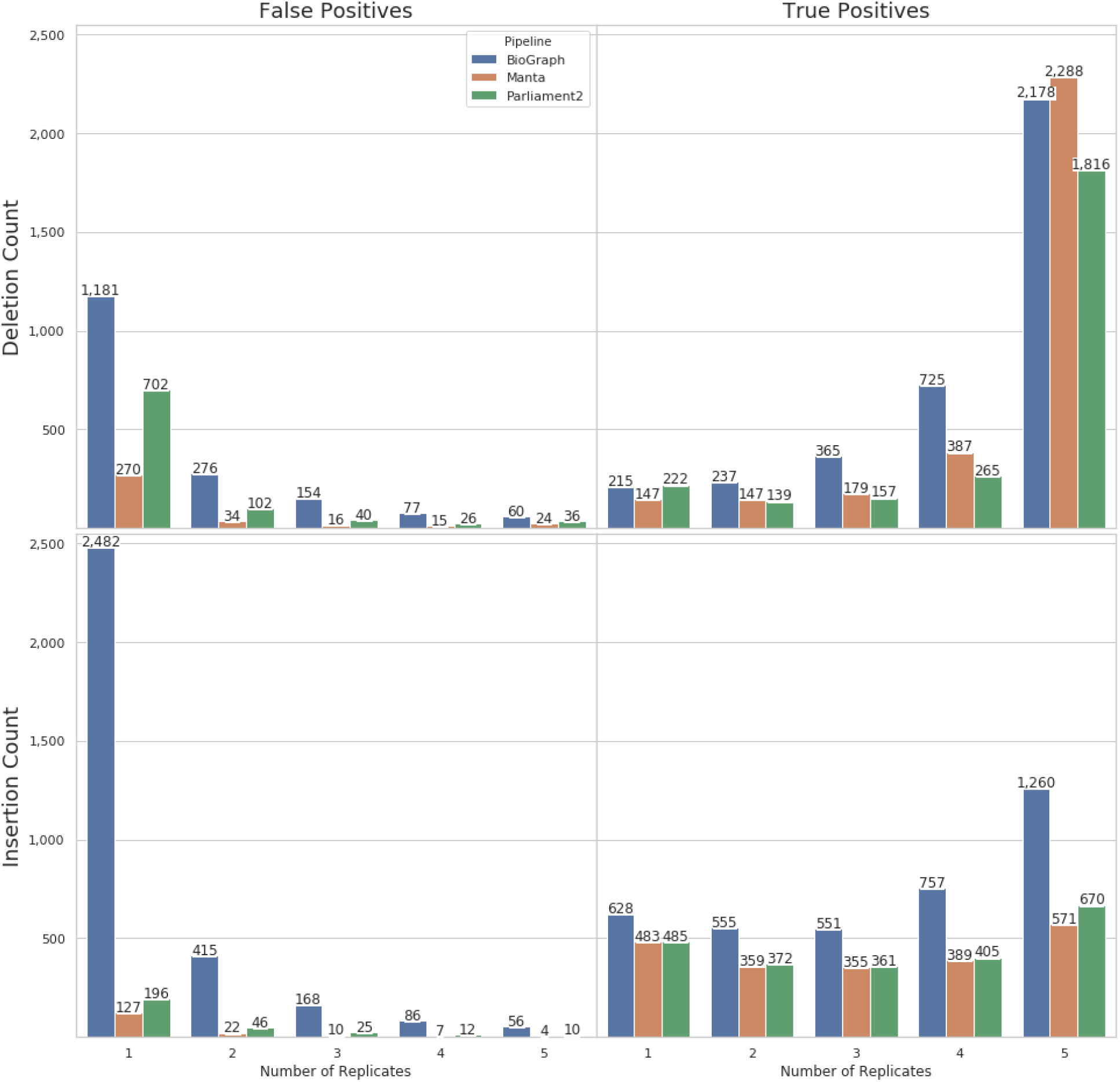
Pipeline consistency of calls across the 5 replicates for each pipeline by false positives and true positives (columns) and by SV type (rows).

Each pipeline assigns a quality score to their calls to quantify the confidence in the variant. These quality scores are useful for controlling the false-discovery rate (FDR). Figure 2 plots a curve of pipeline sensitivity as a function of FDR. By raising the minimum quality score of accepted variants, Biograph can achieve a lower FDR than Manta and Parliament2 while maintaining a higher sensitivity for the same FDR. For example, filtering BioGraph’s calls by raising the minimum acceptable quality score to 50 to achieve a similar FDR returns an average per-replicate FDR of .08 and sensitivity of 48%, compared to Manta’s non-filtered FDR of ≈.09 and sensitivity of ≈43%

**Figure 2.**
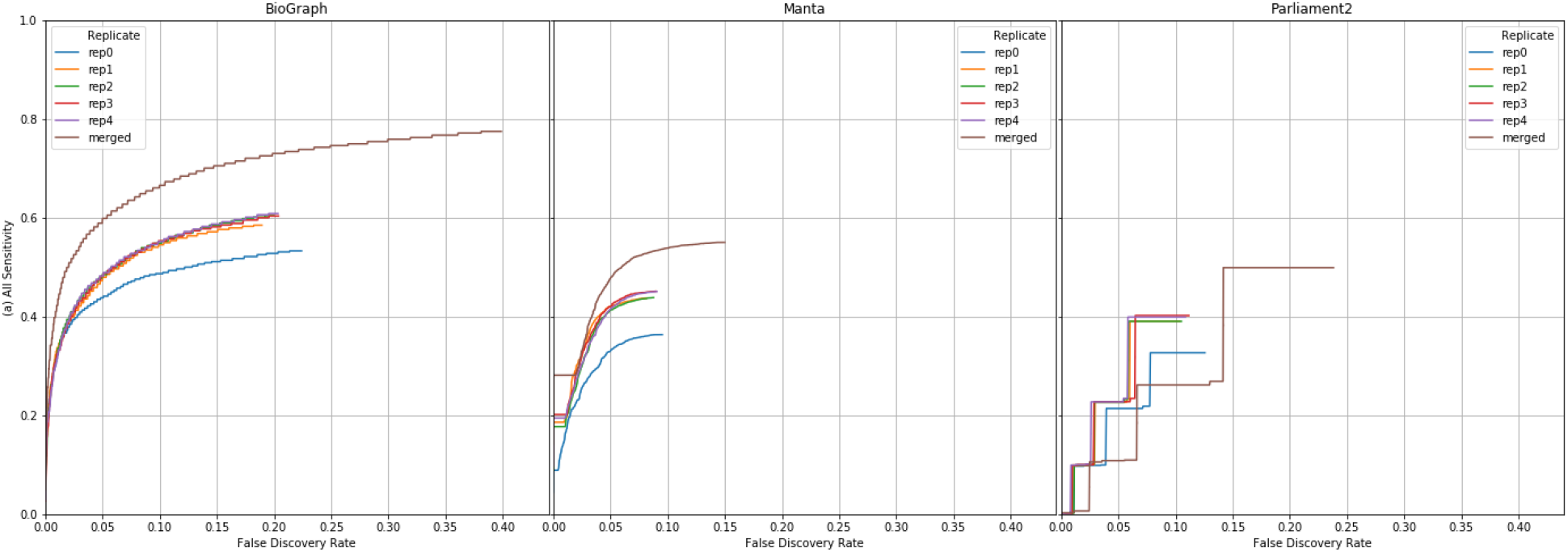
Curves of the sensitivity as a function of FDR for replicates’ calls sorted by quality score. Note approximately 40% of Parliament2 calls are insertions, all of which were reported as having a quality score of zero.

The calls were consolidated between replicates within each pipeline using ‘bcftools merge’ to create a merged VCF. When multiple replicates found identical calls, the highest reported quality score is preserved for the variant (‘merged’ line in Figure 2). Supplementary 4 shows the quality score density distributions of calls in merged VCFs separated by TP and FP (false positive) for each pipeline. These merged quality score distributions range from 28 to 100 (mean 63, median 62) for BioGraph, 20 to 999 (mean 766, median 922) for Manta, and 0 to 13 (mean 4, median 6) for Parliament2.

To assess the cumulative recall from each pipeline across all replicates, we again ran Truvari with the merged VCFs against the GIAB Tier1 SVs (Figure 3a). The BioGraph pipeline discovered 77.5% of GIAB Tier1 SVs in at least one replicate compared to 55% and 51.7% discovered by Manta and Parliament2, respectively.

**Figure 3.**
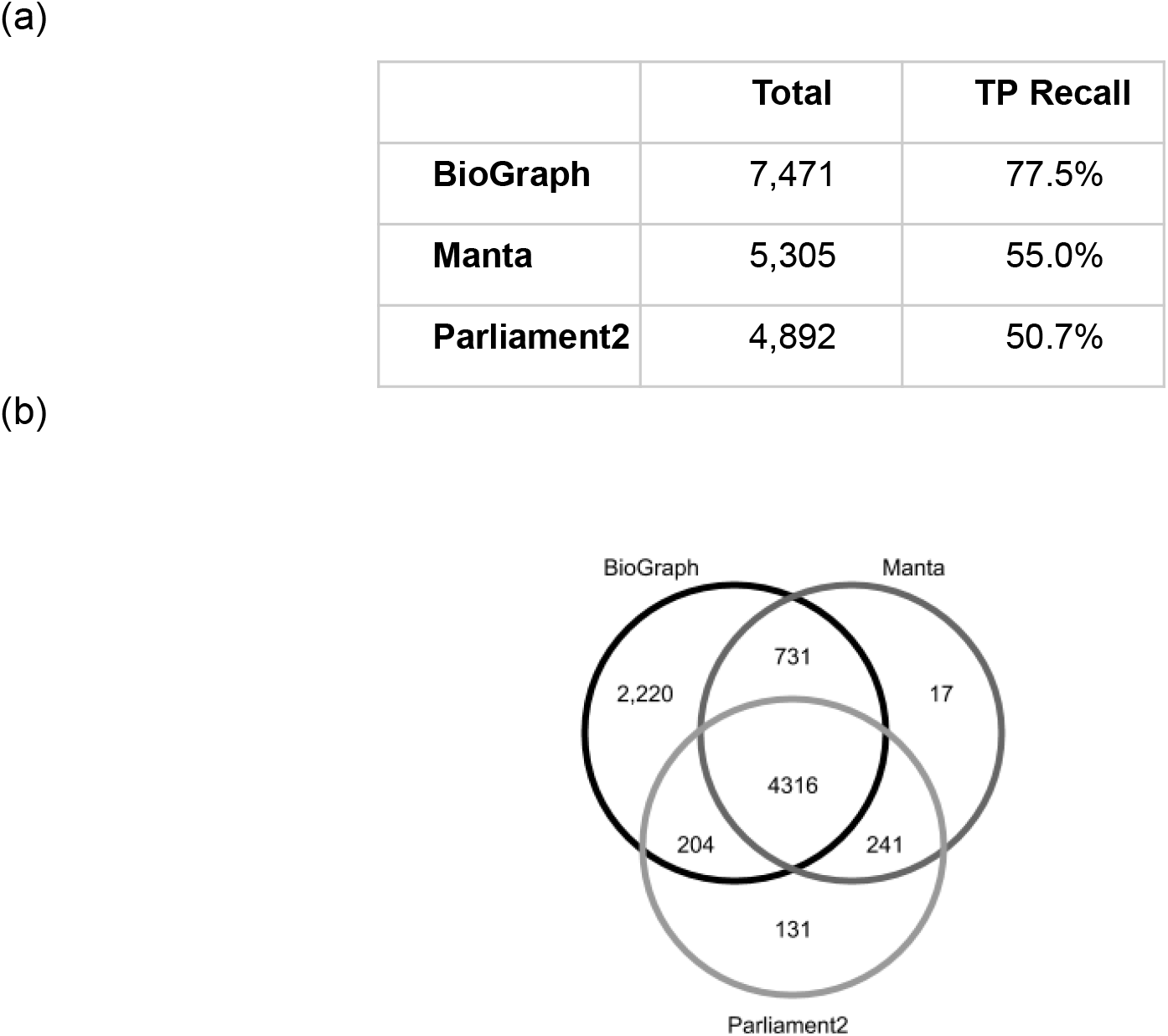
(**a**) Cumulative recall of GIAB TPs from pipelines’ merged project-level VCF. (**b**) Inter-pipeline true-positive Venn diagram. Further details can be observed in Supplementary 5.

The BioGraph pipeline was designed with the goal of discovering as much variation across type and size categories as possible from a single short-read sequencing data experiment. Intersecting the cumulative recall set of GIAB Tier1 variants discovered by each pipeline shows 2,220 of the SVs in the GIAB Tier1 set are only found using the BioGraph pipeline; conversely 389 events found by either Manta or Parliament2 are not called with BioGraph Discovery (Figure 3b). This leaves 1,781 (18.5%) of GIAB Tier1 SVs undiscovered by any of the short-read pipelines used here.

The alleles in the merged VCF from the BioGraph pipeline are used to extend the GRCh37 reference as described in the BioGraph Coverage methods below. Replicates are then queried for coverage against this set of reference variation to create a squared-off project-level VCF (pVCF). We observe that almost all of the true positives discovered using BioGraph Discovery in any one replicate have some read coverage for each replicate (Figure 4). The replicate with the least total coverage (replicate 0), had approximately 20% of deletions with a coverage less than 10x.

**Figure 4.**
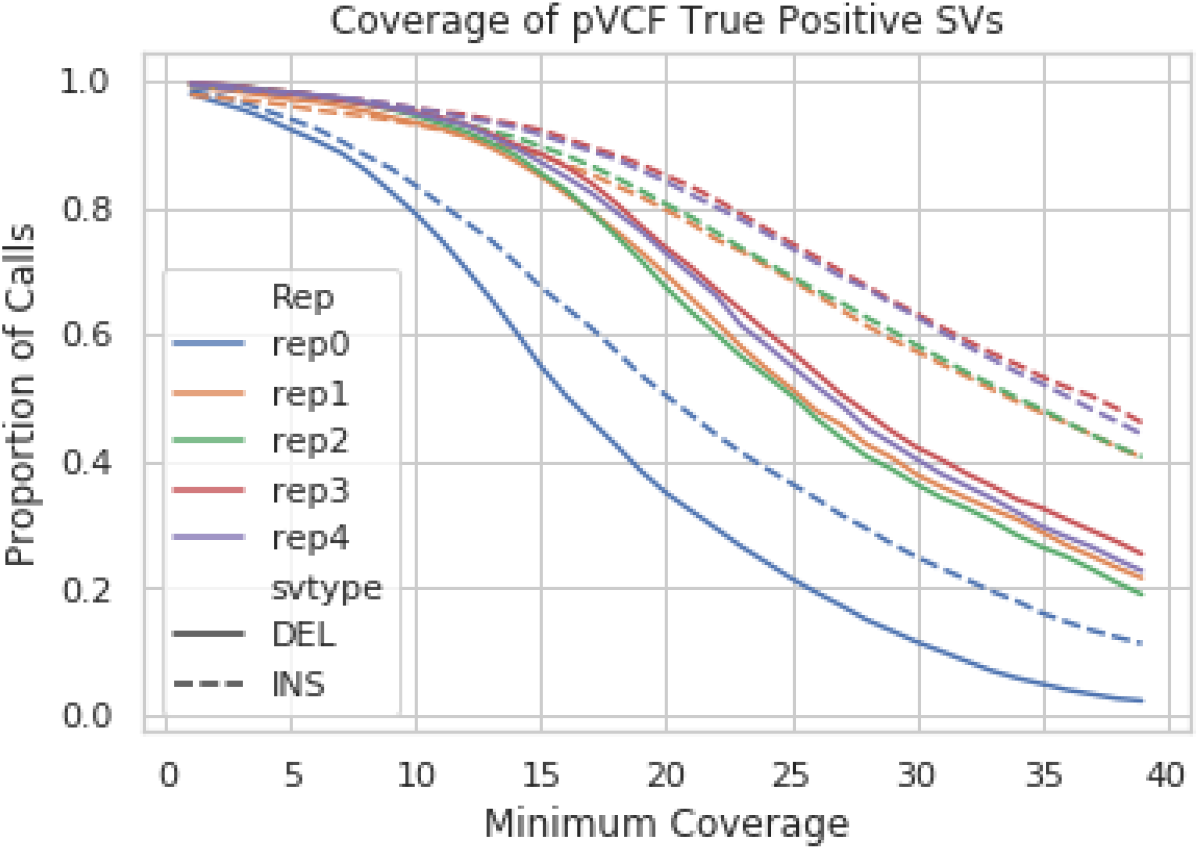
The proportion of the union of true positive variants discovered across replicates using the BioGraph variant calling pipeline (N = 7,471) recovered using BioGraph Coverage with a certain minimum coverage over the variant by replicate number and structural variant type.

Of the 7,471 true positives discovered in at least 1 replicate, 3,438 true positive variants were discovered prior to squaring-off with at least 1x coverage in all 5 replicates (Figure 1). However, by squaring-off coverage for those 7,471 variants in the pVCF, 6,714 true positives had at least 5x coverage in each of the five replicates using BioGraph Coverage (Figure 5). This is an increase from 46% to 90% in the proportion of true positive calls recovered in all five replicates over all true positives represented in the pVCF. Approximately 97% of true positives in the BioGraph produced pVCF were recovered in at least 4 of 5 individuals. We observe in replicate 0 that there were 71 true positive calls discovered independently within the replicate using the BioGraph pipeline having less than 5x coverage. However, leveraging the full set of true positives recovered across replicates with the BioGraph Coverage pVCF, there are 753 variants recovered in replicate 0 where coverage was less than 5x.

**Figure 5.**
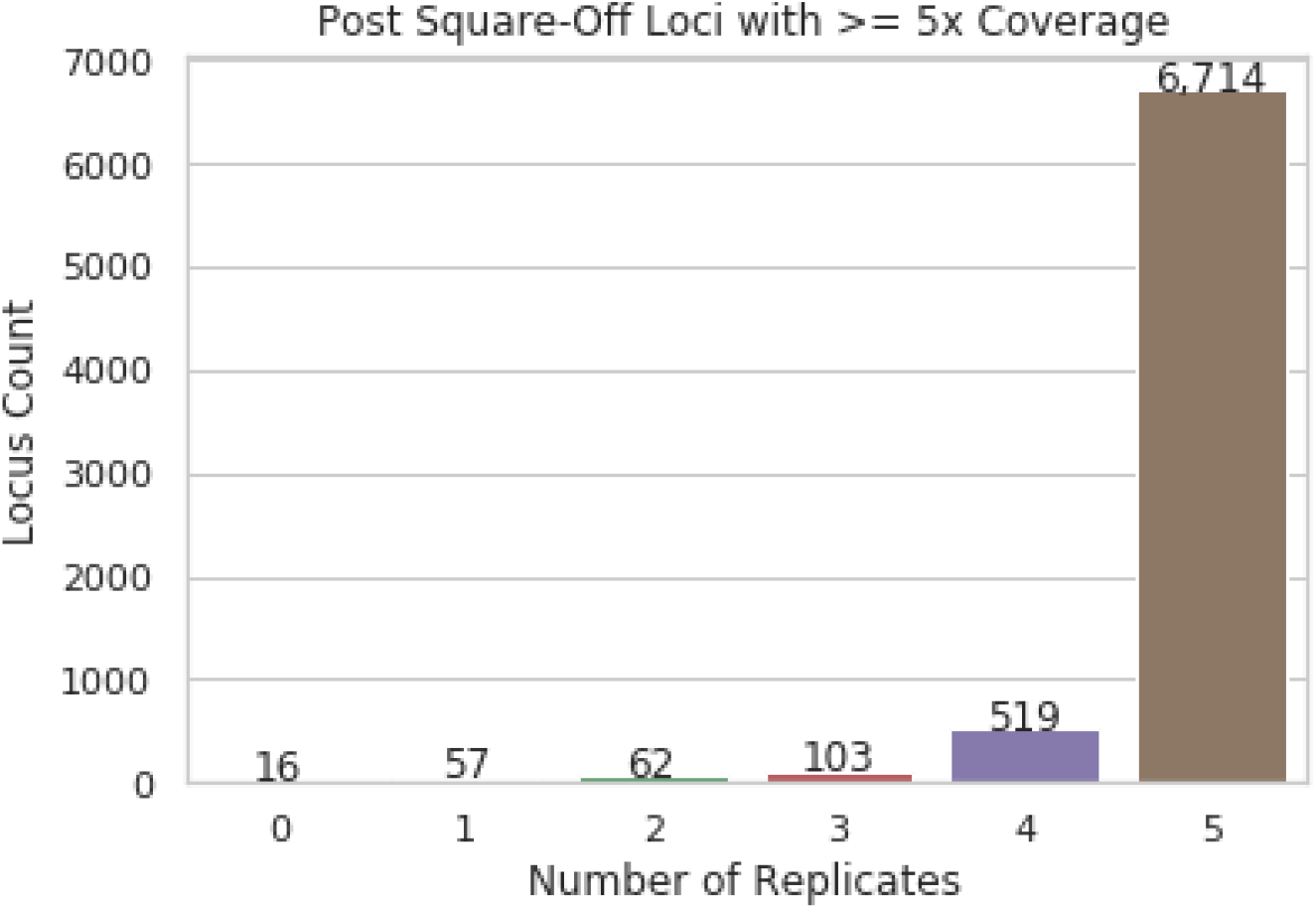
The number true positive variants in the pVCF split into categories indicating the number of replicates in that the variant was recovered with at least 5x coverage using the BioGraph Coverage (N = 7,471).

Within each replicate, the recall of the GIAB truth set increased from an average of 59% per-replicate to 76.9% when including calls recovered using the pVCF and BioGraph Coverage (further details in Supplementary 6). This squaring-off process also increased the consistency at which GIAB truth set alleles were recovered in all five replicates to 69.6%. BioGraph Coverage works by searching for paired reads that match the exact sequence of the structural variant as it is represented in the reference variant graph. If there is a discrepancy in nucleotide sequence in the graph and the nucleotide sequence in the reads then BioGraph Coverage will not report any coverage over the variant. Therefore, this result suggests that approximately 76% of SVs in the GIAB truth set were called using the BioGraph variant calling pipeline with the allelic sequence that matched the exact nucleotide sequence for the variant as represented in the reads.

## Discussion

We leveraged novel data-structures and algorithms to produce a set of structural variants for a well characterized individual. By running BioGraph and other common short-read SV detection pipelines across multiple sequencing replicates, we were able to compare the effectiveness of three different approaches.

Given the ability to rapidly search for reads by sequence using this novel data-structure, we utilized a read assembly method to call variants. Because BioGraph Discovery reports the exact nucleotide sequence and breakpoints of variants, recovery of those variants in other sequencing experiments is possible when there’s less supporting coverage. This suggests that in applying our approach to a larger number of individuals, it may be possible to call an SV in one individual and then recover it in another where that allele is present but there may have been insufficient coverage for discovery, thus leveraging multiple samples to increase the recall per-individual and consistency of variants within a population.

Although we use the Genome in a Bottle set as the primary reference set of variants for benchmarking SVs, another study has published an independently derived set of high quality SVs in the same HG002 individual [11]. As a post-hoc analysis, we used Truvari to determine whether any of the so-called GIAB relative false positives from the three pipelines were found to be true positives against this call set. An average of 611 false positives (≈42% of false positives per-replicate) from BioGraph were found in this orthogonal ‘truth set’. For Manta and Parliament2, on average 60 and 118 calls matched, corresponding to ≈15% and ≈25% of their false positive calls respectively (Supplementary 7). If these false positives are in fact real and were included as part of the base set of variation against which we benchmark, we would see an increase in precision and recall for each pipeline. This suggests that an investigation such as the one conducted in this paper may only provide an approximation of performance that is highly dependent on the completeness of the baseline ‘truth set’. Future work will explore the effects of different properties of these baselines such as sequencing technologies (e.g. short-reads vs long-reads) and variant calling algorithms can have on BioGraph’s relative performance.

## Methods

### Sequencing data

To benchmark, we used five replicates of the Personal Genome Project’s Ashkenazi Jewish Proband HG002. The first replicate is ≈30x sampling of the Illumina HiSeq 150bp paired-end reads from the Genome in a Bottle Consortium [22].

Separately, cell lines from the HG002 individual were sequenced to ≈46X at the Baylor Human Genome Sequencing Center as four technical replicates using the Illumina NovaSeq platform.

### Bioinformatics

The replicates were aligned with BWA [14] and SVs were called with Manta [9] Parliament2 (Zarate). The variant calls from each of the methods were compared to the GIAB v0.6 Tier1 truth set for SV calls (Supplementary 9) using Truvari (Table 1). After the reads were converted to the BioGraph Format, SVs were called using BioGraph Discovery - a whole read overlap reference guided assembly. Then, coverage metrics were generated using BioGraph Coverage - a method to calculate coverage over variant sequences using a dynamic graph reference. Finally, a machine learning classification model, QUALClassifier, was applied to the resulting variant calls with coverage metrics to generate a quality score that can be used to improve precision. The per-replicate set of variant calls in VCF format were then merged and re-run with BioGraph coverage to create a squared-off project-level VCF (pVCF). Steps of the BioGraph pipeline are described in detail below. Computational performance metrics of BioGraph are detailed in Supplementary 1.

### The BioGraph Format

The BioGraph Format (Diagram 1) stores information on one or more genomic samples, and the sequences of bases contained therein. The set of all sequences present in any sample is contained in the Sequence Set (“SeqSet”), and information on which sequences represent full-length reads from a sample is contained in a Read Mapping (“ReadMap”). The BioGraph SeqSet is a BWT-like structure which holds a highly compressed and indexed representation of all possible sequences and allows ‘existential’ queries for the presence/absence of any sequence contained. Additionally, push and drop operations on queries allow traversal between sequences, logically replicating a De Bruijn graph. These properties enable efficient reference-agnostic querying. The result of a SeqSet query is a ‘context’ holding the start/end span over the BWT-like structure with the sequence it represents. This context’s start/end can then be hashed to return information from a ReadMap about which reads begin with the current context’s sequence, the reads’ length, sequencing orientation (direct/complement), and any associated read-pairs’ index in the ReadMap. A reverse hash operation from a ReadMap index to a SeqSet context allows recovery of a read’s full sequence. For details on the BioGraph Format implementation, see [23].

On intake of reads, a read correction method is used to reduce graph complexity. The read correction step reduces sequencing errors with virtually no impact on the ultimate set of variant calls. It works without the need to consider quality scores, and thus requires no base quality score recalibration step. The process is completely reference-free, eliminating the possibility of reference bias [24].

### BioGraph Discovery

The read-indexing applied by the BioGraph Format allows for sequencing experiments to be rapidly queried for the existence of sequences. It also allows for the rapid traversal of overlapping sequences across putative assembly paths. BioGraph Discovery (Diagram 2) leverages these capabilities to perform reference guided assembly, mapping, and variant calling. We begin by walking a query sequence - typically a chromosome from a genomic reference - one base at a time from start to end. At positions of the query sequence where the nucleotide sequence of the read assembly diverges from the query sequence (i.e. a base other than the query’s next base can be pushed onto the assembly sequence) is noted as the start of a potential alternate assembly. That assembly path is extended using the global set of reads contained within the BioGraph Format that have at least 70% read-length overlap between adjacent reads. The assembly is extended in this way until it converges back to the query sequence or once 100,000 bases are assembled without convergence to the query sequence, halting further assembly. This allows allelic sequences to be assembled without the need for bases that overlap the query, which can include large structural variations with lengths greater than that of a single read.

Any assembled paths that diverged from but do not converge back to the query sequence are passed to a second step. In this step, BioGraph Discovery takes those preassembled paths and limits the search space for sequences that can extend the assembly path from all possible overlapping sequences to only sequences from reads where the pair of that read is already within the current assembly or sequences from reads that map exactly downstream of the original breakpoint for that assembly in the query sequence.

All assembly paths are then scored according to the size of the variant, the number of reads comprising an assembly, and whether the read-pairs of reads within an assembly matched back to the query location near the points of divergence/convergence. The paths with the highest scores are added to the list of potential assemblies for each region of the query. This process is then repeated with the reverse complement of the query. Finally, assemblies are deduplicated and the top scoring variants are reported at each locus, up to the expected ploidy of the organism. The result is a Variant Call Format (VCF) file containing SNVs, indels, and structural variants. The variant calls from BioGraph Discovery alone, when compared to the GIAB set shows high sensitivity with low precision (Supplementary 8)

### BioGraph Coverage

Given a flat reference genome, and a set of sequence resolved alleles in a VCF/BCF format, we can create a reference variant graph (RVG). This RVG is an in-memory extension of a given flat reference with paths (branches) that diverge from the reference for each of these variants, creating a graph representation similar to that described in depth in [12]. Edges of the graph represent inter-base reference locations between alleles, and nodes represent nucleotide sequences. The sequences may contain reference bases or alternate alleles. Variants from any number of individuals may be added to the RVG. As SNVs and indels within read length of structural variant calls may be in phase along the reads that support a structural variant, SNVs and indels are contained within the RVG and genotyped as well. We consider this graph to be dynamic as it can be updated with additional alleles from a VCF/BCF file without the need to recompute a global index of the graph.

The BioGraph Coverage (Diagram 3) algorithm walks the graph and searches a BioGraph Format file for read-pairs with concordantly exact matches to the sequences represented. That is, the nucleotide sequence of the read(s) and its pair match exactly and the pair-distance is within the specified insert-size maximum. For each variant, the coverage metrics (e.g. minimum depth over an alternate allele, number of reads spanning an edge) are recorded in an output VCF. This method requires that for a variant to be recalled, the nucleotide sequence of reads in the sample that are associated with the variant exactly match the nucleotide sequence of a path through the graph.

### BioGraph QUALclassifier

The exhaustive, global search for assembly paths relative to a reference performed by BioGraph Discovery finds a large number of false positive assemblies. To reduce the presence of false positives we sought to assign quality scores to each putative allele. A model was created that assigned a quality score for each allele called from BioGraph Discovery based on coverage metrics provided by BioGraph Coverage on those calls. Random forest is an ensemble learning method composed of multiple decision trees, each of which is built from a bootstrap sample of training data using a randomly selected subset of variables. The scikit-learn [25] package’s implementation of RandomForestClassifier was employed in this study. This model was trained on the called alleles from six randomly downsampled coverage titrations (100x, 70x, 50x, 30x, 20x, and 10x) of publically available 300x Illumina data on individual HG002 [22], with each allele’s true positive or false positive state assigned based on the comparison of haplotype-resolved assemblies of long-reads for HG002 [11] against the GIAB v0.6 Tier 1 set using Turvari.

To obtain an unbiased and optimal set of features, three selection methods were proposed to rank the features: feature importance based on gini index, permutation importance and mutual information. The overlapped features chosen by three different approaches resulted in 36 features. The final classification model was trained over 500 trees using 36 coverage metrics and put through a 10-fold cross-validation. The model assigns a quality score (QUAL) to each call, which is the percentage predicted probability for the call to be a true positive. A threshold is used to filter calls (default 28.4). Calls with a quality score below the threshold are removed from the final VCF to reduce the number of likely false positive calls reported. The default threshold was optimized for precision, however it can be adjusted for desired precision-recall performance. The curves in Figure 4 are a useful tool to visualize the sensitivity-specificity tradeoff of the classifier.

### Truvari - software to compare calls to the truth set

To compare structural variant calls to assess performance, we created the tool Truvari [26]. This tool compares a base and comparison VCF file to determine which structural variants match. By default, Truvari matches entries that satisfy the following conditions: calls’ start/end span is within 500bp (refdist); calls have at least 70% size similarity (pctsize); the alternate allele’s haplotype over the events’ spans has at least a 70% Levenshtein-distance ratio (pctsim) - a proxy for sequence similarity. The only non-default matching parameters used were allowing for multiple comparison calls to multiple base calls, and including only calls with a FILTER of PASS in the VCFs. Only events within the GIAB Tier1 high-confidence regions are used when calculating precision and recall.

### BioGraph Coverage across multi-samples’ calls

Since BioGraph Coverage creates a reference variant graph from alleles in a VCF file together with the flat reference over which those variants were called, it is possible to merge variants discovered in multiple samples to create a merged VCF reference variant graph. Many pipelines that consolidate SVs between multiple samples account for the imprecision of breakpoints by using merge heuristics such as variants’ distance from each other or reciprocal overlap between those at the same locus but across samples [10]. Because the BioGraph variant calling pipeline reports precise breakpoints of normalized, sequence resolved variation, it is possible to merge multiple samples’ calls with the same procedure as that for SNP/INDEL variants such as bcftools merge [27]. Once the multi-sample reference variant graph is created from the merged VCFs and reference, samples’ BioGraph Format can be queried for reads that exactly match all putative haplotypes the VCF’s alleles could reconstruct using BioGraph Coverage. Merging these per-sample coverage query results creates a squared-off project-level pVCF.

## Supplementary Materials

https://docs.google.com/spreadsheets/d/1FlRXB_NhLX6lIFdCJ-Woq8eJazIJMYsObWsJEXIB9jM/

**Diagram 1.**
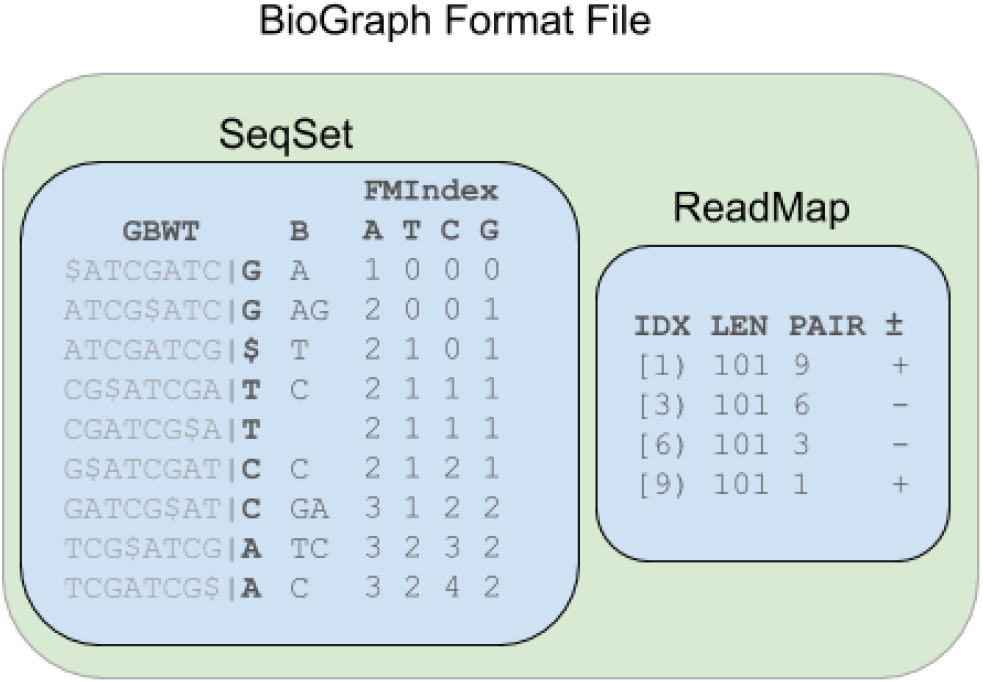
Schematic showing the structure of a BioGraph Format File for a sample.

**Diagram 2.**
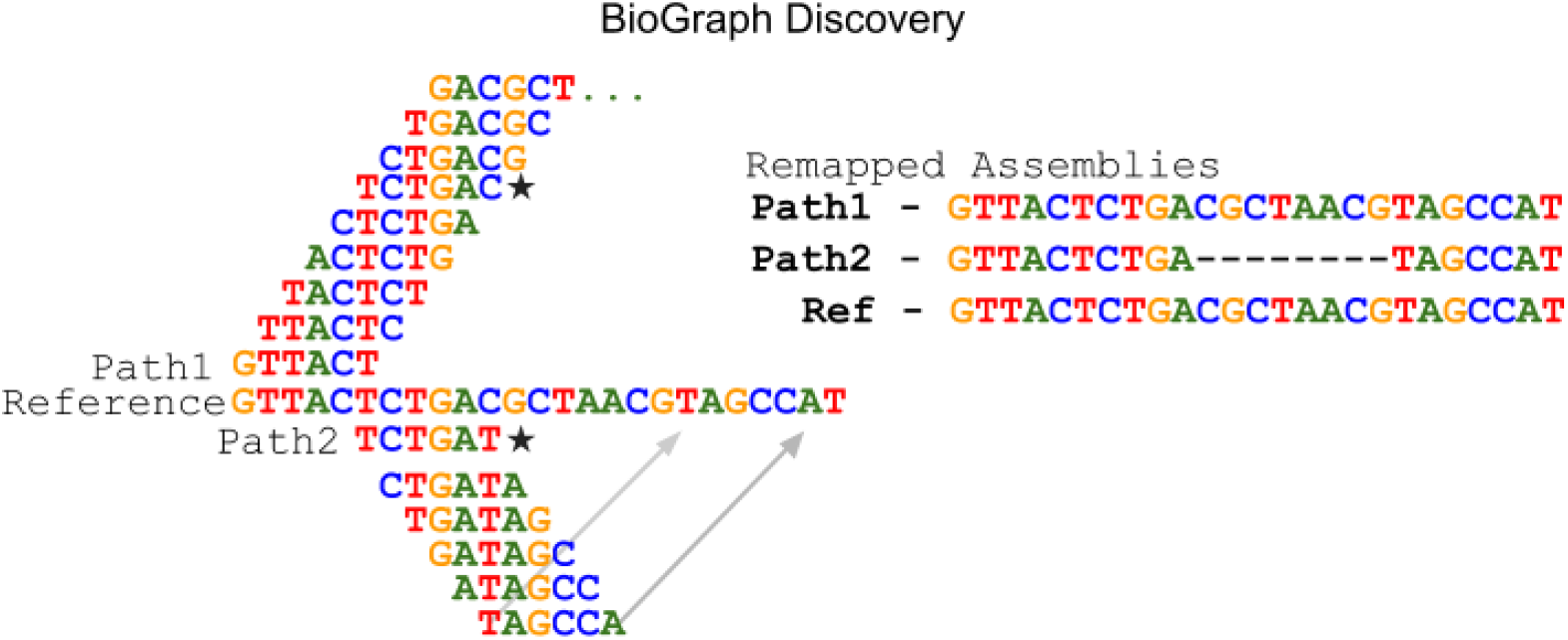
The BioGraph Discovery method using a read length of 6 to discover two alternative alleles.

**Diagram 3.**
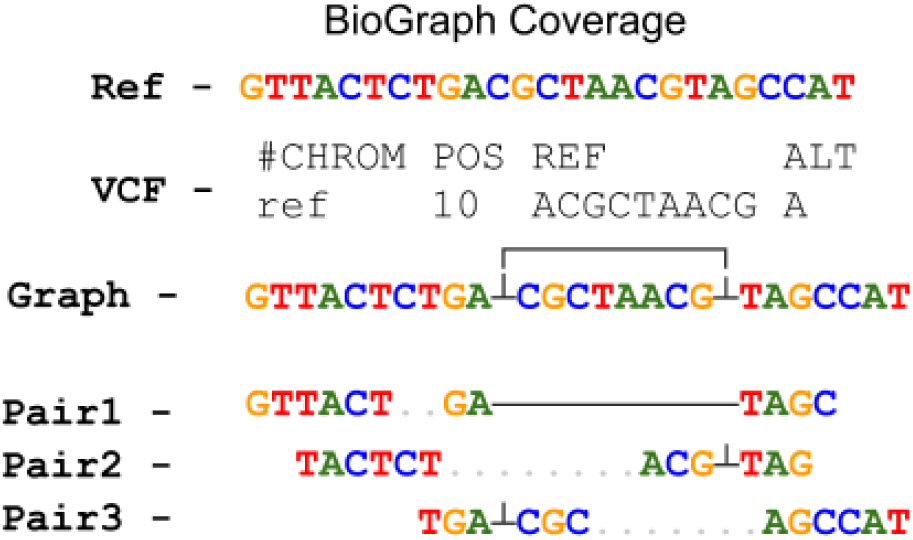
Illustration of how a BioGraph Coverage can map read-pairs to a RVG illustrated with paired reads of 6 nucleotides.

